# Large-scale simulation of biomembranes: bringing realistic kinetics to coarse-grained models

**DOI:** 10.1101/815571

**Authors:** Mohsen Sadeghi, Frank Noé

**Affiliations:** Department of Mathematics and Computer Science, Freie Universität Berlin, Arnimallee 6, 14195 Berlin, Germany

## Abstract

Biomembranes are two-dimensional assemblies of phospholipids that are only a few nanometres thick, but form micrometer-sized structures vital to cellular function. Explicit modelling of biologically relevant membrane systems is computationally expensive, especially when the large number of solvent particles and slow membrane kinetics are taken into account. While highly coarse-grained solvent-free models are available to study equilibrium behaviour of membranes, their efficiency comes at the cost of sacrificing realistic kinetics, and thereby the ability to predict pathways and mechanisms of membrane processes. Here, we present a framework for integrating coarse-grained membrane models with anisotropic stochastic dynamics and continuum-based hydrodynamics, allowing us to simulate large biomembrane systems with realistic kinetics at low computational cost. This paves the way for whole-cell simulations that still include nanometer/nanosecond spatiotemporal resolutions. As a demonstration, we obtain and verify fluctuation spectrum of a full-sized human red blood cell in a 150-milliseconds-long single trajectory. We show how the kinetic effects of different cytoplasmic viscosities can be studied with such a simulation, with predictions that agree with single-cell experimental observations.

---

Lipid bilayers are essential structural elements of living cells, and are important for various cellular functions, such as exo/endocytosis and signal transduction [1, 2]. The unusual properties of biomembranes - they are two-dimensional fluids, but have an out-of-plane elasticity similar to solid sheets - have been the subject of numerous biophysical studies during the past decades [3–7]. Nonetheless, simulating biologically relevant membrane systems remains a challenge [8]. Considering length- and time-scales involved in biological processes, coarse-graining has become an essential approach in membrane simulations [9–12]. Interacting particle reaction-dynamics (iPRD) models are extremely coarse-grained models that have been used to simulate a wide variety of cellular signalling pathways [13–21]. In such simulations, systems containing proteins, lipids, and metabolites are modelled with particles large enough to represent whole proteins or protein domains. Previously, we have proposed a membrane model that seamlessly integrates into iPRD simulations, while accurately reproducing bending rigidity, area compressibility, and in-plane fluidity of biomembranes [22].

While many coarse-grained membrane models may reproduce correct equilibrium properties [10, 23–25], there is currently no satisfactory treatment of the membrane kinetics. Correct kinetics are essential in order to make predictions about not only how fast, but also by which mechanisms biological processes happen: considering that biological functions are far from thermodynamic equilibrium [26, 27], the pathways by which systems relax to steady states or transition between them directly depends on the kinetics. As examples, consider passive transport driven by density fluctuations [28], dynamics of membrane scission by ESCRT proteins [29], as well as dynamin superfamily [30], and questions such as whether vesicle exo/endocytosis rather proceeds by fusion and recycling or by a faster partial fusion (kiss and run) mechanism [31, 32]. Membrane kinetics is thus indispensable when investigating membrane-mediated interactions, binding/unbinding events, and membrane remodelling processes. Also, considering that novel experimental techniques such as dynamic optical displacement spectroscopy (DODS) have made it possible to look at membrane fluctuations resolved at 20 nm and 10 μs range [33, 34], modelling tools that are up to the task of combining the large-scale dynamics of the membranes, while resolving these microscopic scales are needed more than ever.

As membrane-solvent coupling is important for correct membrane kinetics, the issue is more pronounced in the so-called solvent-free membrane models, in which the interactions of membrane particles are adjusted, so as to implicitly account for the missing solvent [11, 35–38]. Discarding solvent particles drastically reduces the computational cost, but leads to unrealistically fast kinetics. A simple correction is the so-called time-mapping [39], to artificially scale the time in order to match the experimental timscale of a specific dynamic property, such as lipid diffusion [24, 40]. But this approach fails when multiple timescales are present and can therefore not improve our ability to predict mechanisms, as it preserves the relative order between timescales.

An alternative is to replace all-atom solvent by a simplified coarse-grained explicit solvent [9, 10, 41, 42], or to use the lattice Boltzmann method to couple particle motions to a grid-based numerical solution of fluid dynamics [43, 44]. Both approaches limit the accessible time- and length-scales, due to additional computations necessary for the fluid response.

Considering both the need for reliable kinetics, and the wide range of scales, a desirable solution for a solvent-free membrane model would be to implicitly incorporate the two major timescales corresponding to in-plane and out-of-plane motions. For this purpose, we propose to use anisotropic stochastic dynamics containing hydrodynamic interactions. Yet, instead of relying on *ad hoc* descriptions of friction and diffusion, or *a posteriori* time-mapping, we derive the governing dynamics of membranes coupled with fluid environments from first principles. As a result, the expected kinetics of the membrane suspended in a solvent is accurately reproduced. Employing this approach, we use our recently developed mesoscopic membrane model and derive dispersion relations of large membrane patches near thermal equilibrium. Our results are consistent with the predictions of a comprehensive continuum-based model. Furthermore, we demonstrate how the incorporated range of hydrodynamic interactions mainly affects the behaviour far from equilibrium, as well as the equilibration rates. Finally, to demonstrate the efficiency and robustness of the proposed method in accessing unprecedented large-scale dynamics in a realistic biological setup, we simulate a human red blood cell and obtain ~150 ms single-trajectories. We successfully compare the power spectral density of red blood cell’s thickness fluctuations with experimental observations based on phase imaging microscopy. We also show the sensitivity of the kinetics to the cytoplasmic viscosity, with potential applications in single-cell profiling and diagnosis.

## I. RESULTS

### A. Anisotropic stochastic dynamics for coarse-grained membrane models

Fig. 1a shows a generic coarse-grained model of the bilayer membrane in which the two leaflets are resolved. In the stable bilayer phase, the particles comprising the membrane diffuse laterally in the membrane domain, while they are coupled to the hydrodynamics of the solvent domain in the out-of-plane direction. The equations of motion of these particles can be written down in a very general form, using anisotropic over-damped Langevin dynamics with hydrodynamic interactions [45, 46],

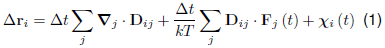

where subscripts *i*, *j* are particle indices, **D**_*ij*_ is the diffusion tensor, **F**_*j*_ is the sum of forces acting on the *j*-th particle, **∇**_*j*_ denotes the divergence with respect to the coordinates of the *j*-th particle, *k* is the Boltzmann constant, and *T* is the temperature. The noise term, ***χ***_*i*_ (*t*), is the outcome of a Gaussian process described by the moments,

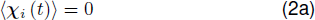

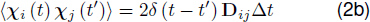

**Figure 1:**
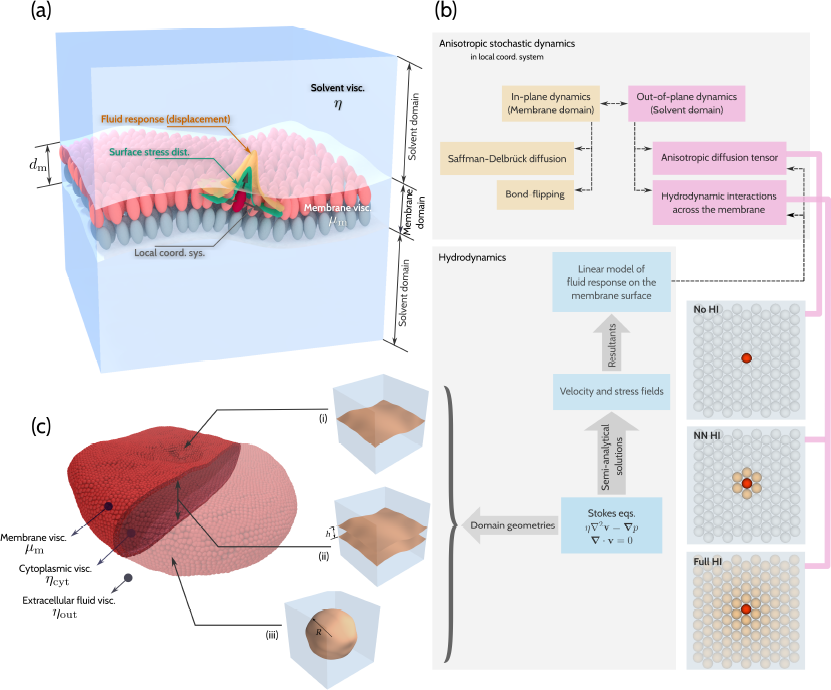
Overview of the proposed method for anisotropic stochastic dynamics of a coarse-grained membrane model **(a)**schematic of a coarse-grained membrane model with the two leaflets resolved. For a membrane suspended in solvent, distinct membrane and solvent domains are designated. The local coordinate system describing the in-plane and out-of-plane directions, as well as the distributed boundary conditions and the fluid response are shown for a selected particle. **(b)**schematic of the proposed method for handling anisotropic stochastic dynamics. **(c)**three distinct idealized geometries used in the derivation of the fluid response: (i) single planar membrane, (ii) parallel planar membranes, (iii) spherical vesicle. The cell membrane of a human red blood cell can be considered to experience fluid responses approximated by a combination of the three geometries.

Several approximations of the **D**_*ij*_ tensor exist for the case of spherical particles floating freely in solvents. Starting from the Stokes-Einstein model, 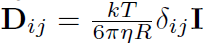[47, 48] (with *η* being the viscosity of the solvent and *R* the particle radius) to the more sophisticated models such as the Oseen [49], Rotne-Prager [50] and Rotne-Prager-Yamakawa [51] tensors. These models are constructed based on analytic solutions to continuum hydrodynamics, albeit with simplifying assumptions, and can be used to include hydrodynamic interactions between particles. But if we consider such a description for densely-packed particles forming a membrane, which usually partitions the space into interior and exterior regions, and through which the solvent cannot easily permeate, it is obvious that a new solution is required.

We construct our anisotropic description via considering a local orthonormal basis at the outer surface of one of the membrane leaflets, and decomposing the displacement of each particle as the sum of in-plane and out-of-plane contributions (Fig. 1a). Thus, the inplane dynamics is dictated by the viscosity of the bilayer membrane, whereas the out-of-plane dynamics involves forces generated due to membrane elasticity and the dissipation through the fluid domain in which the membrane is suspended. These two viscosities differ by 2-3 orders of magnitude [52–54]. As a result, simulation schemes operating on one time-scale cannot reproduce the kinetics efficiently. The main ideas for building an efficient and accurate model for the anisotropic dynamics of membrane particles are:

- The main contribution of solvent-mediated hydrodynamic forces is along the membrane normal. Laterally, the in-plane viscose forces dominate over shearing contributions from the solvent.
- While in-plane hydrodynamics of bilayer membranes can also be studied rigorously [55, 56], a highly coarse-grained membrane model, would benefit little from it. Also, it has been shown that if only large undulation modes are considered, as it is the case with a highly coarse-grained model, the dissipation in the fluid domain dominates over its in-plane counterpart [57].
- For in-plane diffusion in a membrane crowded with proteins, the hydrodynamics simplifies to a collision-based dynamics, resulting in a Stokes-Einstein-like diffusion [58].

Based on these arguments, we propose the following form for the diffusion tensor,

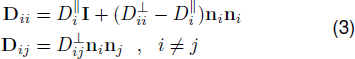

where *D*^‖^ and *D*^⊥^ respectively represent the in-plane and the out-of-plane diffusion coefficients and **n**_*i*_ is the unit vector normal to the membrane surface at the position of the *i*-th particle. Provided that the values of *D*^‖^ and *D*^⊥^ are known for all the particles forming the membrane, we can make use of Eq. (1) to efficiently obtain particle trajectories. As the breakdown chart in Fig. 1b suggests, we can use the Saffman-Delbrück model of the diffusion of cylindrical inclusions in fluid sheets to obtain the in-plane components, 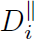 [59, 60],

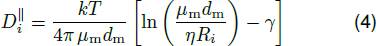

where *R*_*i*_ is the radius of a cylindrical inclusion in the membrane domain, *μ*_m_ and *d*_m_ are the viscosity and thickness of the membrane domain, *η* is the viscosity of the surrounding medium, and *γ* ≈ 0.577 is the Euler–Mascheroni constant. Unfortunately, there are no readily available description of hydrodynamics yielding out-of-plane components, 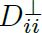 and 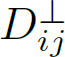. To derive numerical values for these components, we have developed semi-analytical solutions to the Stokes equations in select idealized geometries (see Figs. 1b and 1c) [61]. The underlying approach is to use Gaussian distributed velocity or stress boundary conditions as the test input, and numerically integrate the resulting fields over selected patches on the surface of the membrane, i.e. find the generated forces and displacements which describe the fluid response (Fig. 1a). For the simple case of a single planar membrane, this procedure yields a closed form solution for 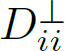[61]

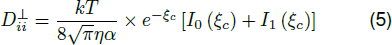

where 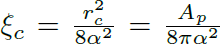 with *α* being a scaling parameter corresponding to the width of the Gaussians used as input, *A*_*p*_ is the area per particle, and *I*_0_ and *I*_1_ are the modified Bessel functions of the first kind. Unfortunately, such closed-form expressions are not available for arbitrary geometries. However, based on the solutions presented in [61], it is possible to find numerical results specific to a membrane model, using the scheme depicted in Fig. 2a. An exemplar outcome of such calculation, for a specific membrane model [22] in different geometries, are compiled in Fig. 2b.

Comparing values of 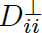 with the diffusion coefficient given by the Stokes-Einstein formula, and 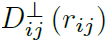 with the magnitude of hydrodynamic interactions predicted by the Oseen tensor (Fig. 2b), demonstrates the short-coming of both descriptions when applied to membrane particles. Interestingly, compared to free-floating spherical particles described by the Stokes-Einstein model, membrane particles have a much higher mobility in the out-of-plane direction. This could be explained by noting that the solvent only affects the particles in the membrane normal direction. As we have demonstrated in [61], for a single planar membrane, the Oseen tensor under-predicts the magnitude of hydrodynamic interactions at least by a factor of 2.

**Figure 2:**
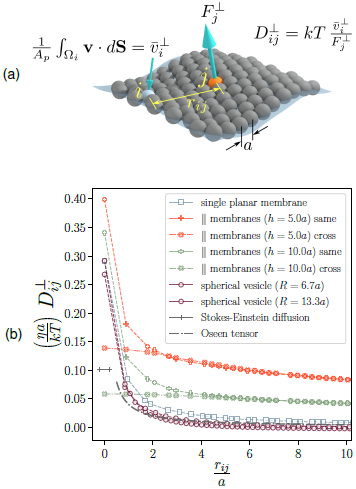
Numerical calculation of the out-of-plane component of the diffusion tensor, 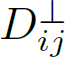. (**a**) definition of the 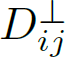 based on 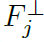, the force existing between the fluid domain and particle *j*, and the corresponding effective out-of-plane velocity, 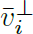, attributed to particle *i*. The effective velocity results from averaging the velocity field of the fluid, in the direction normal to the membrane, in the vicinity of the membrane particles. (**b**) compilation of numerical values of the 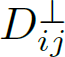 as a function of in-plane inter-particle distance, *r*_*ij*_. Results are respectively given for a single planar membrane, two sets of parallel membranes with the given inter-plane separations, and two spherical vesicles with the given radii. In the case of parallel membranes, the in-plane distance is measured solely based on planar coordinates, and for spherical vesicles, it is defined along the geodesics. For these calculations, the parameter *α* is chosen equal to 0.1*a*. Values given by the Stokes-Einstein relation, as well as the Oseen tensor are shown for comparison.

Comparing the results for different geometries given in Fig. 2b, two conclusions are in order:

- Generally speaking, membrane curvature has little effect on hydrodynamic interactions. This can be readily observed in Fig. 2 by comparing the results corresponding to single planar membrane, and the two spherical vesicles with different radii. The deviation is only important for 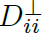 components.
- There is a significant difference between the mobilities and hydrodynamic interactions of parallel membranes compared to single membrane patches, with generally larger 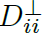 and 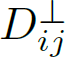 values. Also, hydrodynamic interaction between particles on the opposite sides are non-negligible. This increased mobility can be considered as the cause for fluctuation-magnification observed for membranes near walls [62]. The closer the membranes are together, the more pronounced are the confinement effects.

Finally, one important aspect of the proposed method, which drastically distinguishes it from previous descriptions such as the Oseen tensor, is that the hydrodynamic interactions between particles are calculated in the presence of the rest of the particle system in a given geometry. While in more generic methods, due to the complexities arising from recurrent interactions between particle pairs, triplets, etc. enhancing the approximation to include such effects is immensely difficult.

### B. Near-equilibrium kinetics of a planar membrane patch suspended in aqueous solvent

Based on the diffusion tensor introduced in Sec. I A, we investigate the predicted kinetics of square-shaped planar membranes patches coupled to an aqueous solvent, when they fluctuate near their equilibrium configuration. Here, we employ the membrane model introduced in [22], where the bilayer consists of closely-packed laterally mobile particle dimers (see Methods section III A for details), but other coarse-grained membrane models [9–12, 63, 64] could be used as well. It is well-known that hydrodynamic interactions are generally long-ranged [65], but it is worth noting that in contrast to systems of free particles, the hydrodynamic interactions between membrane particles have to compete with large forces resulting from the bending rigidity of the membrane. We thus expect the solvent effects to manifest mostly as local dissipation, and only to a lesser degree as force transmission across the membrane. To further study this, we consider three different models of hydrodynamic interactions (Fig. 1b):

1. **No HI**: No hydrodynamic interactions exist between membrane particles. The only non-zero diffusion coefficients are 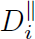 and 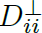, whose values are correspondingly taken from Eqs. (4, 5). This results in a diagonal diffusion matrix and uncorrelated local anisotropic dynamics.
2. **NN HI**: Hydrodynamic interactions are only included between nearest-neighbour particles that are defined by construction of our membrane model, resulting in a fast implementation. To introduce correlated random displacements resulting from these interactions, we construct local diffusion matrices and use Cholesky decomposition to transform a vector of independent normally distributed random variables to a correlated vector obeying Eq. (2b).
3. **Full HI**: Hydrodynamic interactions are implemented across the membrane, with global diffusion matrices compiled in each iteration. To avoid the 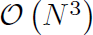 cost of a full Cholesky decomposition, we use the approach developed by Geyer and Winter to approximate the correlated random forces [66].

We have used a forcefield developed with the same approach as in [22], with parameters given in Methods section III A. We also consider aqueous solvent with viscosity *η* = 0.890 mPas (Details in Methods section III D). Based on the chosen forcefield and diffusion tensors, we found a timestep of 0.5 ns to produce stable trajectories. Compared with timesteps of at most 20 ps for the same model without hydrodynamics [22], there is an apparent 25-fold increase in performance. Though the cost of integrating stochastic dynamics in each model of hydrodynamics is to be considered. As we will show, with the cheap **No HI** model, the speed-up significantly enhances the timescales available to exploration with the current model, pushing simulation times to the 100 ms range.

The kinetics of the membrane patch in the vicinity of thermodynamic equilibrium can be determined by looking at how fast its thermally-induced undulations relax. This mode-dependent relaxation dynamics yields the so-called dispersion relation of the membrane. A reliable theoretical description of the dispersion relations of the membrane, that has been shown to be consistent with experiments, is given by the continuum model of Seifert et al. In this model, the two membrane leaflets are resolved, and height fluctuations of the membrane are coupled to the hydrodynamics of the solvent as well as the variations of lipid densities inside its leaflets. This model predicts the relaxation dynamics of the out-of-plane motion of each undulatory mode, designated by the wave vector **q**, to follow a biexponential decay (See Sec. III B for details). The theoretical values for the two corresponding frequencies, denoted here as 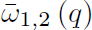, depend on the viscosity of the solvent, *η*, as well as the bending rigidity of the membrane, *κ*, area compressibility modulus of one leaflet, *K*_area_, membrane thickness, *d*_m_, and a phenomenological inter-leaflet friction coefficient, *b* [3, 67, 68]. The asymptotic values of these two frequencies for the range of wave vectors investigated here are 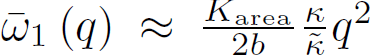 and 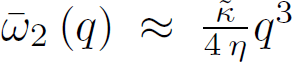, which respectively signify the slow and fast dissipation regimes [41]. The slow dissipation regime (the slipping mode) corresponds to in-plane density fluctuations, as well as the friction between the two leaflets, while the fast regime (hydrodynamic mode) is due to the viscous loss in the fluid. Importantly, the slow mode depends strongly on the internal dynamics of the membrane, and thus, on the details of the coarse-grained model, while the fast mode is only a function of the emergent bending rigidity and the prescribed hydrodynamics of the solvent, making it our focus of interest here.

In each case, we have used two different values of the scaling factor *α*, equal to 0.01*a* and 0.1*a*, to obtain diffusion tensors. The value of *α* relates the cut-off radius defined in terms of the lattice parameter of the model, *a*, to the hydrodynamic length-scale arising from the chosen width of the Gaussian boundary conditions [61]. We have performed simulations of membranes starting from a flat initial state, with respective trajectory lengths of 1.5 ms for the **No HI**, and 1.0 ms for **NN HI** and **Full HI** cases. The reason for this choice becomes clear when we look at far-from-equilibrium dynamics. As we are interested in near-equilibrium kinetics here, we discard an initial portion of each trajectory, allowing for complete equilibration. The kinetics in this initial segment of trajectories, as well as the required length of it, will be the focus of the next section.

For each sampled frame, the height function, *h* (*x*, *y*), is obtained by mapping the vertical position of particles to a regular grid (Fig. 3a). Fast Fourier transform is used to obtain values of *h*_**q**_ (*t*), which are used to calculate the tion of the ensemble averages and the auto-correlation functions in Eqs. (7a, 7b). As a result, we obtain dispersion relations by fitting biexponential functions to the autocorrelations (Figs. 3b-3d) and power spectra of the equilibrium thermal undulations (Figs. 3e-3g).

**Figure 3:**
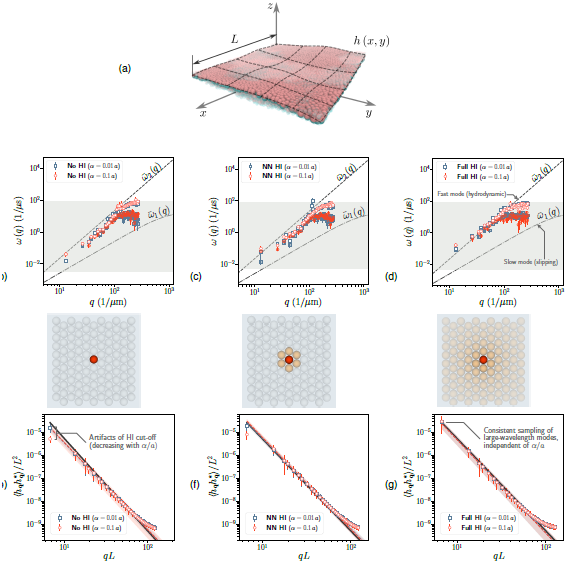
Simulating thermal undulations of a membrane patch suspended in water. (**a**) an overlay of several simulation snapshots performed using the mesoscopic membrane model with the lattice parameter of *a* = 10 nm. Also shown is a schematic depiction of the smooth height function, *h* (*x*, *y*) fitted to particle positions in each frame. **(b) - (d)** dispersion relations for the membrane patch, calculated based on the assumption that the fluctuations in the amplitude corresponding to a wave vector **q** relax biexponentially with frequencies *ω*_1,2_(*q*). The fast regimes (higher *ω*) are depicted with empty symbols, while filled symbols are used for the slow regimes. Results are shown for two different choices of the scaling factor *α*/*a*. Predictions of the continuum model based on the Helfrich functional coupled with compressible leaflets (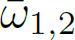 functions from Eq. (7b)) are included for comparison. The light grey region signifies the range of frequencies available, depending on the sampling rate and the length of the trajectory. The three cases correspond respectively to the **No HI**, **NN HI**, and **Full HI**assumptions regarding the hydrodynamic interactions. **(e) - (g)** power spectra of thermal undulations of the membrane patches in the aforementioned simulations. Dashed lines are fits of the function *C*(*qL*)^*n*^ to the data, whereas the solid black line is the prediction of the continuum model given by Eq. (7a). Shaded areas denote 99% prediction intervals of the power-law fits.

If we look at the power spectra of thermal undulations, it is obvious that all cases follow the expected equilibrium behaviour given by Eq. (7a) (depicted as the solid black lines in Figs. 3e-3g). This result is reflective of the fact that (a) we have indeed sampled from equilibrium configurations of the membrane, and (b) fluctuation-dissipation theorem holds for the stochastic dynamics developed here. The latter pertains to the validity of the general form of the diffusion tensor (Eq. (3)) as well as the correlated noise terms being correctly approximated in the simulations (Eq. (2b)). A closer inspection reveals the presence of small deviations for long wavelength modes in **No HI** and **NN HI** cases. These deviations are artefacts of hydrodynamic interaction cut-off. We suspect this to be due to the rotational dynamics of the normal vectors, which are not completely described when hydrodynamic interactions are cut short. When the hydrodynamic effects are present across the membrane (the **Full HI** case), rotational dynamics are naturally included. These deviations decrease with *α*/*a*, and almost vanish for *α*/*a* = 0.01. This is to be expected, as when *α*/*a* is small, the cut-off radius in terms of *a*, or number of particles included in the hydrodynamic interactions, translates to a larger radius in terms of the actual length scale of the hydrodynamics of the solvent, reducing the artefacts.

To use Eqs. (8) and obtain the theoretical eigenvalues 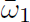 and 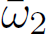, the viscosity of the solvent, as well as thickness, viscosity, bending rigidity, and area compressibility modulus of the membrane, are all a priori given values used in the parametrization of the model (see Sec.III A in Methods). Only the inter-leaflet friction, *b*, is unknown. Experimental determination of *b* is rather difficult, and is based on measuring the velocity difference between the two leaflets in pulling tethers from vesicles [69, 70]. The resulting values are in the range 10^8^ to 10^9^ N s m^−3^ [69, 71]. Yet, all-atom or coarse-grained simulations predict much smaller values in the 10^6^ to 10^7^ N s m^−3^ range [70, 72, 73]. It is to be expected that the inter-leaflet friction coefficient be highly sensitive to the resolution with which the lipids are modelled. Here, we have used the value of 10^7^ N s m^−3^ to calculate 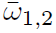. It should be considered as a reference value, which only has significance regarding the slow mode with the frequency of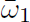.

Comparing simulation results with the expected dispersion relations (Figs. 3b-3d) show that in all cases, very good approximations of the desired fast (hydrodynamic) mode are achieved. Interestingly, the slow mode is also present, and within the range expected based on the parameters fed to the model, in spite of our *ad hoc* choice of the inter-leaflet friction. The reproduction of fast mode, with the asymptiotic behaviour of 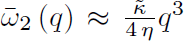, proves that the framework laid out here is indeed capable of yielding trajectories with the realistic kinetics that is expected from membrane elasticity coupled to hydrodynamic effects. The choice of the hydrodynamic scale parameter, *α*, at least in the range of inspection, has little effect on the simulated dispersion relations. This will prove helpful in robust application of this framework to other membrane models of different scales. While the **Full HI** case best reproduces the fast mode, even without the inclusion of hydrodynamic interactions, the viscous dissipation is captured rather well in the **No HI** case. This allows for efficient near-equilibrium sampling of membrane dynamics with coarse-grained models, relying solely on a locally anisotropic stochastic dynamics.

### C. Far-from-equilibrium kinetics of the suspended membrane patch

We studied the equilibrium behaviour of membrane patches in aqueous solvents in Sec. I B, and showed both the equilibrium distribution as well as kinetics in the vicinity of equilibrium are correctly reproduced by the method developed here. Now, we can also investigate the dynamics of the same system, at states far from thermodynamic equilibrium.

We consider the path that each system takes towards equilibrium. The initial flat configuration of the membrane is a state far from thermodynamic equilibrium. To quantify how fast membrane patches evolve from this state towards equilibrium, we consider the slowest process, which is the evolution of the longest undulatory wavelength, **q**_0_. We look at how the quantity 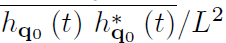, which corresponds to the energy of this mode, evolves with time. Arguably, there exists no theoretical description similar to Eq. (7b) to describe this non-equilibrium evolution. Thus, we use a generic exponential expression in the form of 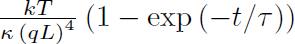 and use the parameter *τ* as a relaxation time (Figs. 4a and 4b).

**Figure 4:**
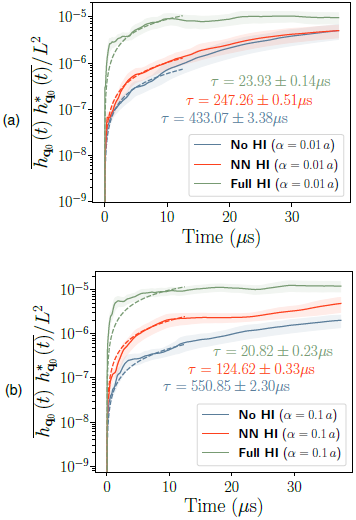
Relaxation of the energy of the largest undulation mode, from a state far from equilibrium to the equilibrium value. Results are given for all the hydrodynamic cases for (**a**) *α* = 0.01 *a* and (**b**) *α* = 0.1 *a*. Dashed lines are fits of the function 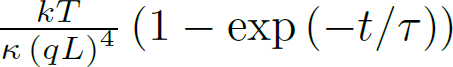 to the data. Corresponding values of the timescale *τ* are given in each case.

It can be seen that the range to which the hydrodynamic interactions are present clearly affects the nonequilibrium behaviour. The **No HI** case has the slowest, and the **Full HI** the fastest dynamics towards equilibrium. If we compare the values of *τ* to the theoretical timescale of fluctuations of the same mode about the equilibrium configuration, which is 20.11 μs based on the model described in Sec. III B, the difference between the near-equilibrium and far-from-equilibrium dynamics becomes apparent. Only for the **Full HI** case the relaxation times are comparable. Also, in this case, the choice of *α/a* has little effect on the dynamics and timescales are consistent for both cases.

For states far from thermodynamic equilibrium, inclusion of the hydrodynamic interactions with the largest possible range ensures a quick descent to the minimum free energy state. But the small fluctuations about the equilibrium state are equivalently well modelled with different hydrodynamic interaction ranges.

### D. Fluctuation profile of a human red blood cell

To demonstrate our membrane dynamics framework on a complex, biophysically relevant example, we consider a human red blood cell. Aside from being the classical subject of membrane biophysics studies [74–76], there has been a rather recent interest in profiling individual red blood cells for diagnostic purposes [77–82]. Diseases such as hereditary spherocytosis and sickle cell, that change the mechanics of the red blood cell membrane as well as the rheology of its cytosol, can be diagnosed by looking at the fluctuations of individual red blood cells using phase-shift microscopy techniques [78]. Small changes in the haemoglobin concentration of red blood cells can also be detected with the same approach. Another example is the pathology of the malaria disease, which is closely related to the mechanical response of the red blood cell to the invasion of parasite’s merozoites [83–85]. Finally, the simple picture of the red blood cell thermal “flicker” has also recently been revised by showing that there is ATP-dependent active mechanisms, possibly reorganizing the spectrin cytoskeletal structure, leading to deviations from the expected equilibrium picture for timescales beyond 100 ms [33, 86–88].

Here we use our membrane model, in conjunction with the hydrodynamics approach introduced and verified on membrane patches, to investigate the thickness fluctuations of the red blood cell. We have performed simulations using a 3D model of the cell obtained from refractive index tomography. Further details of the simulation setup are laid out in methods section III C. Due to the double-layer nature of the membrane model, we were able to assign different solvent hydrodynamics coupled to the two sides of the membrane, i.e. we have used different dynamics for the leaflets facing the inside or the outside of the cell. The former is considered a surface in contact with a large body of aqueous solvent, while the latter is modelled as parallel membranes with their inner space filled with the more viscous cytoplasmic matrix. Equipped with the method described in Sec. I A, we are able to simulate red blood cells with different internal viscosities. Our sole aim here is to show how different kinetics affects observables such as the magnitude of cell vibrations.

We simulated the red blood cell for as long as ~150 ms to obtain reliable statistics as well as large-scale kinetics. The baseline viscosity for the cytosol is taken to be *η*_cyt_ = 4.5 mPa s [88]. We consider the thickness profile of the red blood cell lying in the *xy*-plane to be given as *d*_RBC_ (*x, y*), and obtain mean and variance of the thickness of the RBC, 〈*d*_RBC_〉 and 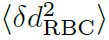, by sampling over the surface of the cell in each frame. As is seen in Fig. 5b, the simulated distribution of the mean thickness over the cell’s surface, quantitatively matches the phase-shift imaging data. We have applied fast Fourier transform of the thickness variance time series, using the well-established Welch’s method [89], to obtain power spectral density (PSD) of the thickness fluctuations(Fig. 5c). The dashed line in the figure shows the least-square fit of a power law to these data points, which gives the exponent of 1.33 ± 0.03. A similar power law has been observed for healthy human RBC’s [74, 87]. Simple theoretical models based on small amplitude fluctuations yield an exponent of 5/3 [90], while Brochard and Lennon’s classic experiments put the exponent in the range 1.45 to 1.3 for *f* > 1 Hz [74]. Using the phase imaging data provided by Park et al., we calculated thickness profiles (Fig. 5b) and power spectral densities of the thickness fluctuation in a similar manner. The power law fitted to the simulation results has been extended to pass through the experimental data points (Fig. 5c).

**Figure 5:**
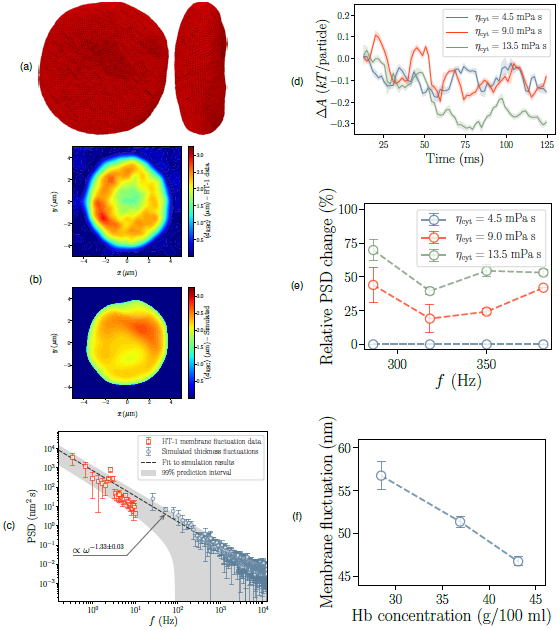
Simulating the dynamics of a human red blood cell, (**a**) top and side views of a snapshot of the red blood cell as modelled with the mesoscopic membrane model. (**b**) mean cell thickness profiles from experimental measurements (holotomography and phase-shift imaging with HT-1, Tomocube Inc., Republic of Korea) compared with the simulated counterpart. (**c**) comparing power spectral density (PSD) of thickness fluctuations of the red blood cell between simulation and HT-1 experimental data. The power-law is fitted to the simulation results and extended to the experimental low-frequency range for the sake of comparison. (**d**) free energy change during the first 125 ms of simulations as a function of time, given for three different choices of the viscosity of the cytoplasm. (**e**) relative change in the PSD for the given viscosities, using the lowest viscosity of *η* = 4.5 mPas as the baseline. (**f**) magnitude of membrane fluctuations versus the haemoglobin concentration of the blood cell cytosol.

A very good agreement can be observed between the two, especially considering the envelope formed by the prediction interval for this power law, based on the uncertainty in the simulation data points.

We have repeated this simulation for two other values of the cytoplasmic viscosity, namely, *η*_cyt_ = 9.0 mPas and 13.5 mPas. We have calculated the non-equilibrium evolution of the Helmholtz free energy of red blood cells with the three different internal viscosities (Fig. 5d). Details of free energy estimation are given in methods section III C. As expected, the free energy moves towards its minimum value with time, but it can be seen that the path taken in the free energy landscape is significantly different for the two lower viscosity cases compared with the high-viscosity one. This result points to the fact that different kinetics can push the system along different configurational pathways, potentially altering path-dependent phenomena.

Finally, to demonstrate how the dynamical framework developed here can pertain to quantities important in diagnosis, we have compared the values of power spectral density corresponding to a select range of frequency, when the cytoplasmic viscosity is increased from the baseline value *η*_cyt_ = 4.5 mPa s. The relative PSD change is measured compared to the values corresponding to this baseline (Fig. 5e). The significant change in PSD for the higher viscosity cases, which is easily measurable based on the fluctuation spectrum, shows how the kinetics of the membrane affects a global mechanical behaviour of the cell. In case of the red blood cell, this could point to an anomaly in a diseased cell. To illustrate this point, we compared the maximum value of membrane fluctuations pertaining to largest undulatory modes for these three cases, but transformed the cytoplasmic viscosity to the haemoglobin (Hb) concentration of the cytoplasm. We used a simple empirical expression [91] for this conversion. The seemingly linear correlation between the membrane fluctuation and Hb concentration (Fig. 5f) has been previously suggested based on quantitative phase imaging techniques [78]. Such a result, with potential diagnosis applications, would be impossible to achieve without relying on a large-scale dynamical scheme that can produce kinetics based on first-principles and empirically meaningful input parameters.

## II. DISCUSSION

We have introduced a framework for incorporating the dynamics of a bilayer membrane suspended in a fluid environment. With the introduction of a novel diffusion tensor that fully describes anisotropic dissipative as well as hydrodynamic effects, we have implemented a robust method to tackle the two-scale dissipative dynamics governing the particles in contact with the membrane and the solvent domains. Combined with a general over-damped Langevin integrator, this approach allows us to significantly increase the timestep used with our membrane model compared to deterministic integrators, easily achieving hundreds-of-millisecond-long trajectories necessary for equilibration and sampling of slow dynamics in membranes of cellular scale.

To investigate the reproduction of expected kinetics in a highly coarse-grained model, we used our particle-based mesoscopic membrane model to obtain dispersion relations for planar membrane patches. Making use of the physical properties used for parametrization of the membrane model, we successfully compared the numerically calculated relaxation dynamics with the predictions of a comprehensive continuum model, incorporating three distinct dissipation modes: out-of-plane viscous loss, in-plane density fluctuations, and interleaflet friction. Despite the simple construction of the membrane model, the anisotropic stochastic dynamics combined with the realistic description of hydrodynamics has made it capable of mimicking these dissipative modes very well, while reproducing a very good estimate of their respective timescales. As this is the result of the stochastic dynamics based on first principles, and not a model-dependent after-the-fact correction for time scales, it is in general applicable to any coarsegrained membrane model. Note that based on previous coarse-grained simulations, if similar kinetics were to be produced using explicit inclusion of solvent particles, at least 10 solvent particle were to be considered per each membrane particle [92]. Even so, the kinetics could only be investigated at a very small scale, because it is rather expensive to equilibrate even sub-micron-sized membrane patches [93]. Such an explicit solvent approach is out of question when considered on cellular scales, while our proposed implicit solvent scheme achieves the same dynamics with a fraction of the computational cost.

We also investigated the far-from-equilibrium kinetics predicted by this approach. We showed how the range of hydrodynamic interactions affects the relaxation time and the equilibration rate, with much faster approach to equilibrium with larger interaction ranges. These results showcase another benefit of the hydrodynamic method introduced here, namely fast equilibration of large membrane patches. It is conceivable to use the more expensive **Full HI** or **NN HI** approaches to equilibrate the system, while using the cheaper **No HI** scheme for equilibrium sampling.

To showcase the promise of this approach in achieving large-scale simulations in biological systems, we performed a 150-millisecond-long simulation on a particle-based model of the human red blood cell. Our approach allowed using different hydrodynamic descriptions for the interior and exterior of the cell, as well as, different values of the cytoplasmic viscosity. Based on the resulting trajectories, we calculated the thickness fluctuations of the red blood cell and showed its power spectral density to asymptotically match the experimental measurements done via phase imaging microscopy, verifying that the proposed dynamical method can thus be expected to reproduce reliable kinetics in a biological system of realistic proportions. We showed the significant effects due to different kinetics being dictated by different cytoplasmic viscosities, and demonstrated how these different could be useful for diagnosis purposes.

The general form of the diffusion tensor introduced here (Eq. (3)) can be used beyond the idealized geometries for which we gave semi-analytic results (Fig. 2b). Our approach can smoothly be integrated into numerical grid-based methods such as the lattice Boltzmann, to represent progressive linearisation of the fluid response, as the membrane acquires complex geometries. This would introduce a significant computational gain, as the grid-based method only needs to be invoked when a significant change in geometry or environment is detected.

The application of the method laid out here is by no means limited to the given examples. The cases studied here prove the reliability of the dynamical scheme, as well as its applicability in complex simulations. But the general scheme can be tailored to the specifics of many other coarse-grained models. The large-scale simulations made possible by this approach have a wide range of applications in studying non-equilibrium pathways of membrane-related cellular process such as exo/endocytosis. This facilitates end-to-end simulation of these complex processes with a unified approach, especially in non-equilibrium scenarios.

## III. METHODS

### A. Mesoscopic membrane model

The membrane model, used for the simulations presented here, is the same as developed in [22]. The bilayer is modelled as formed by particle-dimers in a close-packed arrangement, with a variable lattice parameter in the range of 10 nm, while the two leaflets are resolved. The force field pertaining bonded interaction is given by the following interaction potentials [22],

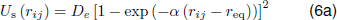

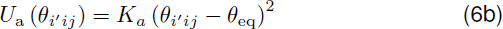

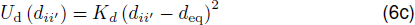

Particles belonging to each leaflet are connected to their nearest-neighbour counterparts via Morse-type bonds (Eq. (6a)). Also, harmonic angle-bending potentials given by Eq. (6b) act against the out-of-plane rotations of these bonds (the primed index designates the opposing particle in a dimer). Finally, particles in a dimer are connected via harmonic bonds of the form described by Eq. (6c), which keeps the two leaflets together. Potential parameters for *U*_s_ and *U*_a_ are obtained using the parameter-space optimization technique described in [22]. The physical properties of the membrane used as input for the simulations in Secs. I B and I C are listed in Tab. I. The values of the potential parameters, obtained or chosen for simulations, are summarized in Tab. II.

**Table I:**
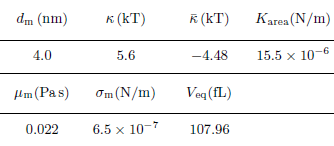
Properties of the membrane used for the parametrization of the membrane model used with simulations in Secs. I B and I C. Values of the bending rigidity, *κ*, and Gaussian curvature modulus, 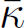, are based on data given in [94–97], while for area compressibility modulus, *K*_area_, data from [98–101] have been considered.

**Table II:**
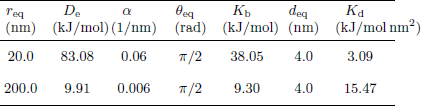
Membrane model force field parameters for the interaction potentials given in Eq. (6), used for simulations in Secs. I B and I C.

### B. Dispersion relations

We take *h*(*x*, *y*) to be a smooth height function fitted to particle positions (see Fig. 3a). Using fast Fourier transform, we find *h*_**q**_ (*t*), which is the amplitude of an undulatory mode with the wave vector **q**. In thermal equilibrium, the ensemble average and the time-dependent relaxation dynamics of these undulatory modes, for a square membrane patch of side *L*, in the absence of any in-plane tensions, are given by following relations [3, 67, 102],

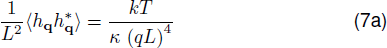

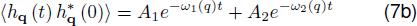

where *κ* is the bending modulus of the membrane. In the continuum-based model of Seifert et al. [3, 67, 102], the relaxation frequencies *ω*_1_ and *ω*_2_, are the eigenvalues of the time evolution operator, −**Γ**(*q*) **E**(*q*), with the following definition,

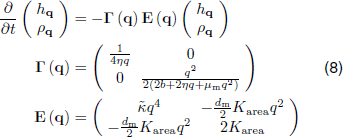

with *K*_area_ being the area compressibility modulus of one leaflet, 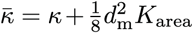 the effective bending modulus, and *b*the inter-leaflet friction coefficient.

### C. Red blood cell

Simulation setup is built using a 3D model of a healthy RBC, obtained through refractive index tomography (courtesy of YongKeun Park [78, 81, 82] and Tomocube Inc., Republic of Korea). The model is captured at a mean resolution of ~100 nm, and consists of a triangular mesh with a wide size distribution. In order to apply the membrane model to this geometry, we have used two reference sets of forcefield parameters, corresponding to lattice parameter values of 20 nm and 200 nm. We have employed an interpolation of the parameters to each bond, based on its neighbouring mesh size. Also, a correction based on the coordination number of particles is locally applied.

The properties of the red blood cell used in developing the model, apart from the elastic constants *κ* and 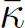, include the viscosity of the membrane domain, *μ*_m_, the cytoplasmic viscosity, *η*_cyt_, pre-existing membrane surface tension, *σ*_m_, the area compressibility modulus, *K*_area_ and the equilibrium volume, *V*_eq_. The surface tension is applied by shifting the equilibrium distances of bonds compared to the lattice parameter dictated by the input mesh. As the membrane undergoes little internal reorganization on the scales of interest due to the presence of cytoskeleton, the bond-flipping Monte Carlo moves, described in [22] to implement laterally liquid membranes, are not used. Volume preservation is enforced through a volumetric potential *U*_υ_ = *K*_vol_ (*V* − *V*_eq_). Values of the input parameters used for parametrization, as well as the resulting forcefield parameters are respectively given in Tabs. III and IV.

**Table III:**
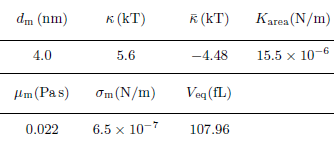
Properties of the red blood cell used for the parametrization of the membrane model used with simulations in Sec. I D [87, 88, 103, 104].

**Table IV:**
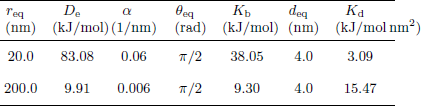
Similar to Tab. II, for red blood cell simulations of Sec. I D.

To calculate the change in the Helmholtz free energy of the red blood cell, Δ*A* = Δ*E* − *T*Δ*S*, estimates of the internal energy, *E*, as well as the entropy, *S* are needed. The internal energy is calculated based on the time averages of the sum of kinetic and potential energies sampled during the simulation. The potential energy is simply the sum of all bonded contributions, while the kinetic energy is estimated based the equipartition theorem. Entropy estimation is done in the Fourier space using the time-dependent probability distribution corresponding to each vibration mode, summed as −*k*Σ_*i*_*p*_*i*_log*p*_*i*_. Other contribution to the entropy from the side walls of the cell, or configurational changes below the resolution of the grid used for the fast Fourier transform, are thus ignored, and are assumed to stay constant for the duration of these calculations.

### D. Simulations

Particle trajectories are obtained by updating the positions of particles according to Eq. (1). The diffusion tensor is updated in each integration step based on instantaneous normal vectors. Normal vectors are calculated for triangles formed between in-plane bonds and averaged for each particle based on its neighbouring triangles. All the simulations are performed at *T* = 298 K and we have chosen water with the viscosity of 0.890 mPa s as the surrounding fluid.

To sample from microstates describing a tensionless membrane, the in-plane degrees of freedom are coupled to the Langevin piston barostat [105]. Application of this barostat has the advantage of seamlessly fitting into the stochastic integrator already used for the in-plane degrees of freedom (Eq. (1)). Thus, the barostat parameters controlling the fluctuation timescale and dissipation of the “piston” are chosen such that they represent a medium similar to the continuation of the membrane patch in the simulation box.

### E. Software

Calculation of diffusion tensors are performed using codes developed in Python, with the help of SciPy computational package. The Python package Matplotlib is used for plotting the results.

Simulations based on the particle-based membrane model [22] are performed using an in-house specific-purpose software. The code is developed using C++. Multithreading parallelization is employed for enhanced performance.

Both codes are available from authors upon request. The software package Visual Molecular Dynamics (VMD) is used for some visualisations [106].

## Conflicts Of Interest

There are no conflicts of interest to declare for this study.

## Acknowledgements

The authors wish to thank YongKeun Park and Geon Kim from department of physics, Korea Advanced Institute of Science and Technology (KAIST) for providing the three dimensional mesh and membrane fluctuation data of the human red blood cell.

This research has been funded by Deutsche Forschungsgemeinschaft (DFG) through grants SFB 958/Project A04 “Spatiotemporal model of neuronal signalling and its regulation by presynaptic membrane scaffolds”, SFB 1114/Project C03 “Multiscale modelling and simulation for spatiotemporal master equations”, and European Research Commission, ERC StG 307494 “pc-Cell”.

## Author Contributions

MS and FN designed research, MS conducted research and developed software. MS and FN wrote the paper.

